# Titer estimation for quality control (TEQC) method: a practical approach for optimal production of protein complexes using the baculovirus expression vector system

**DOI:** 10.1101/258624

**Authors:** Tsuyoshi Imasaki, Sabine Wenzel, Kentaro Yamada, Megan L. Bryant, Yuichiro Takagi

**Affiliations:** Department of Biochemistry and Molecular Biology, Indiana University School of Medicine, 635 Barnhill Drive, Indianapolis, Indiana 46202, USA

## Abstract

The baculovirus expression vector system (BEVS) is becoming the method of choice for expression of many eukaryotic proteins and protein complexes for biochemical, structural and pharmaceutical studies. Significant technological advancement has made generation of recombinant baculoviruses easy, efficient and user-friendly. However, there is a tremendous variability in the amount of proteins made using the BEVS, including different batches of virus made to express the same proteins. Yet, what influences the overall production of proteins or protein complexes remains largely unclear. Many downstream applications, particularly protein structure determination, require purification of large quantities of proteins in a repetitive manner, calling for a reliable experimental set-up to obtain the protein or protein complexes of interest consistently. During our investigation of optimizing the expression of the Mediator Head module, we discovered that the ‘initial infectivity’ was an excellent indicator of overall production of protein complexes. Further, we show that this initial infectivity can be mathematically described as a function of multiplicity of infection (MOI), correlating recombinant protein yield and virus titer. All these findings led us to develop the Titer Estimation for Quality Control (TEQC) method, which enables researchers to estimate initial infectivity, titer/MOI values in a simple and affordable way, and to use these values to quantitatively optimize protein expressions utilizing BEVS in a highly reproducible fashion.

## Introduction

The baculovirus expression vector system (BEVS), introduced about 30 years ago [1–5], has become an essential tool for expression of many eukaryotic proteins and protein complexes in insect cells [6–8]. BEVS has been proven powerful for structure determination of membrane proteins such as GPCRs [9], AMPA receptor [10], NMDA receptor [11], and channel rhodopsin [12], which are inherently difficult to work with, as well as cell surface receptors [13–15]. Moreover, development of the MultiBac system, which is suited for expression of multi-protein complexes, has opened new avenues for structural studies of multi-protein complexes [16–21]. Key technological advancements of the BEVS lie in the ease and efficient generation of recombinant baculoviruses harboring single genes [1, 5], or in case of the MultiBac system, harboring multiple genes, enabling expression of numerous proteins simultaneously [22–24]. Well-established standard procedures for BEVS allow a streamlined workflow, accounting for its ease of use and high adaptability to routine laboratory operations for many users. Despite of all these technological advances to simplify the generation of recombinant baculoviruses, when it comes to protein expression, the key indicator for optimal expression of proteins remains elusive. For example, there is no clear answer to the question of whether or not a user would obtain twice as much protein if twice as much baculovirus is added to the insect cells. The situation is a marked difference from the bacterial expression system in which key factors for optimization of protein expression (e.g. IPTG concentration) have been well established [25].

A lack of clarity regarding optimization of protein expression in BEVS may be attributed to conflicting results from previous studies [26–32]. In these studies, MOI (Multiplicity Of Infection: how many infectious units per cell) was considered to be a key indicator for expression of a model protein, β-galactosidase. While some of these studies showed that varying MOI values had no significant effects [26–28], others showed high MOI [29, 30] or even low MOI [28, 29, 31, 32] conditions resulted in optimal protein expression. The results of these studies did not provide a clear answer about the correlation between MOI and protein production. The problem is a lack of understanding the physiological outcome of MOI values: for example, MOI=2 means having twice as much virus per cell, but how it affects the cells and how it represents protein expression levels are uncertain. Even if MOI is a good indicator, classical titer/MOI measurement methods, such as the plaque assay [33], and the immunohistochemical staining method (Clontech Inc), are too cumbersome and time-consuming to be practically used for high throughput expression studies. Other titer/MOI measurement methods such as qPCR [34], flow cytometric assays (FCA) [35], measurement of cell diameter change of post infected cells [36], and a cell line containing eGFP for titer measurement [37] are less tedious. However, they require specialized equipment, and setting them up is not trivial. Therefore, these methods are not practical for many users, whose goal is to produce proteins of their interest and not to measure MOI per se. Furthermore, the condition at which the titer is measured differs vastly from that at which protein expression is carried out, prompting a question as to how relevant such a measurement is for protein production. Taken all together, a simple and affordable experimental method to quantify and optimize expressions of proteins or protein complexes using BEVS needs to be established for many end users.

Mediator Head module is an essential sub-complex of the Mediator complex, which plays a key role in transcription regulation in eukaryotes [38]. We used BEVS to generate the Mediator Head module for structural and functional studies [16, 39]. We often encountered the problem of inconsistency in expression level of the Head module, which prompted us to investigate a way to optimize expression of the Mediator Head module. Our research led us to discover the ‘initial infectivity’ as an excellent indicator of overall production of protein complexes, and we devised a simple and affordable method to estimate this initial infectivity without the need for any specialized equipment or setting. Our finding led to the development of the ‘Titer Estimation for Quality Control’ (TEQC) method, which enables users to quantify and optimize expression of protein complexes in insect cells, ensuring quality control of protein expression in the BEVS.

## Materials and Methods

### Maintenance of insect cells

Hi5 and Sf9 cells were obtained from Expression Systems (Davis, CA). The cells were maintained in ESF921 media (Expression systems) in shaker flasks at 27°C. Both cells were split to a density of 0.5 × 10^6^ cells/ml every 3 days.

### Construction of baculovirus transfer vectors for expression of multi-protein complexes

All vectors used in this study are summarized in Supplemental Table S1. Construction of the baculovirus transfer vector for expression of modified Mediator Head module used for our structure determination was described in [16]. Construction of the baculovirus transfer vector bearing the genes encoding the yeast Mediator Middle module is as follows: Open reading frames (ORFs) of genes encoding Med7, Med21, Med9, Med4, Med10, Med19 and Med1, were amplified by PCR from the yeast (*S. cerevisiae*) genomic DNA. Med31 with a C-terminal HA and 10xhistidine tag (Med31-HA-10His) was amplified by PCR from pBacPAK9-Med31-HA-10His vector, which was generated by sub-cloning MED31 gene into BamHI and XhoI sites of pBacPAK9 followed by addition of HA and 10xHis tag sequences by two rounds of QuikChange site-directed mutagenesis (Agilent Technologies). The PCR products were cloned into SphI and SmaI sites (MCS1), and BamHI and HindIII sites (MCS2) of the pFL vector using the SLIC method [40] in the following pair-wise combinations: Med7-Med21, Med9-Med4, Med10-Med31-HA-10xHis-tag, and pFL-Med19-Med1, yielding pFL-Med7-Med21 (pYT53), pFL-Med9-Med4 (pYT56), pFL-Med10-Med31-HA-10His (pYT55), and pFL-Med19-Med1 (pYT57), respectively. PmeI and AvrII fragment from pYT53 or pYT55 was sub-cloned into SpeI and NruI sites of pYT56 or pYT57, yielding pFL-Med4-Med7-Med21-Med9 (pYT87), and pFL-Med10-Med19-Med1-Med31HAHis (pYT85). Finally, PmeI and AvrII fragment of pYT87 was sub-cloned into SpeI and NruI vector of pYT85, yielding pFL-Med10-Med4-Med7-Med21-Med9-Med19-Med1-Med31HAHis (pYT90).

Construction of the baculovirus transfer vector bearing the genes encoding the subunits of yeast TFIIF, Tfg1, Tfg2 and Tfg3 is as follows: The C-terminal FLAG and TAP tagged TFG1 gene, yTFG1-FLAG-TAP, was amplified by PCR from pDt/g1g2 vector [41] (gift from Ponticelli at State University of New York). ORFs of genes encoding yeast Tfg2 and Tfg3, were amplified by PCR from the yeast genomic DNA. The PCR products were cloned into SphI and SmaI sites (MCS1), and BamHI and HindIII sites (MCS2) of the pFL vector using the SLIC method [40], yielding pFL-Tfg1-TAP-Tfg2 (TI150). The PCR product for Tfg3 was cloned into BamHI and HindIII sites of pUCDM vector, yielding pUCDM-Tfg3 (pYT154). Fusion of the pTI150 and pYT154 vectors by Cre recombinase (NEB) yielded a transfer vector (TI153), containing all 3 genes encoding the yeast TFIIF. Construction of the baculovirus transfer vector bearing the genes encoding the subunits of yeast core TFIIH, is as follows: ORFs of genes encoding Tfb5, Rad3, Tfb2, Ssl1, Tbf4 and Tfb1-10xHis tagged, were amplified by PCR from pBacPAK9 vectors bearing corresponding genes as described previously [42]. The PCR products were sub-cloned into SphI and SmaI sites (MCS1), and BamHI and HindIII sites (MCS2) of pFL, pUCDM, and pSPL vectors in pair-wise fashion, yielding pFL-Tfb5-Rad3 (TI132), pSPL-Tfb2-Ssl1 (pYT444), and pUCDM-Tfb1-10His-Tfb4 (pYT465), respectively. Fusion of all three vectors by Cre recombinase (NEB) yielded a transfer vector (TI419), containing all 5 genes encoding the yeast core TFIIH. Open reading frames (ORFs) of genes encoding yeast Cycline C (CycC), and CDK8 genes were amplified from the yeast genomic DNA. DNA sequence encoding 10xhistidine tag was added to the C-terminus of CycC gene during PCR. The PCR products were cloned into SphI and SmaI sites (MCS1), and BamHI and HindIII sites (MCS2) of the pUCDM vector using the SLIC method [40], yielding pUCDM-CycC-10His-CDK8 (pYT379). Fusion of pYT379 vector and an empty pFL vector by Cre recombinase (NEB) yielded a transfer vector (TI600) for expression of CDK8-CycC-10xHis complex. Construction of human Taf8-Taf10 expression vector has been previously described [43].

### Virus production and storage

Productions of recombinant baculoviruses in Sf9 cells were performed as described [23]. Liquid viruses were stored at 4°C. Frozen viruses were generated as described (Supplemental protocol) [44]. Frozen virus stocks were stored under liquid nitrogen.

### Estimation of Multiplicity of Infection (eMOI)

The initial infectivity is defined as the infectivity 24 hours after infection (I24). An estimation of the initial infectivity is denoted as eI24. The multiplicity of infection (MOI),which is a ratio of infectious units per cell, can be estimated using the following two equations:

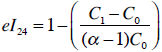

Estimation of MOI = eMOI

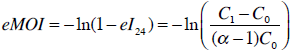

Where a growth constant, α, is the growth rate of uninfected insect cells in a 24 hours period, C_0_ is an initial cell number, and C_1_ is the cell number 24 hours after addition of virus. The detailed procedure for estimation of infectivity and MOI is described in the supplementary protocol in Supplemental Information.

### Titer measurement by Expression Systems, Inc

Our liquid viruses were shipped to Expression Systems (Davis, CA) for titer measurement. Titer measurements were performed via flow cytometric analysis (FCA) for expression of the baculoviral coat protein gp64 by Expression Systems, Inc as described [35]. The numbers of gp64 positive cells identified by FCA were used to calculate the infectivity of the baculovirus expressing the Mediator Head module.

### Expression of the multi-protein complexes in the insect cells

200 ml culture of Hi5 cells (1.0 × 10^6^ cells/ml) or Sf9 cells (1.5 × 10^6^ cells/ml) was infected with the amount of viruses indicated. Infected cells were incubated at 27°C for 96 hours, and were harvested by centrifugation, frozen in liquid nitrogen, and stored as a pellet at −80°C until future use.

### Purification of protein complexes

The Mediator Head module was purified as described previously [45]. The yeast Mediator Middle module, core TFIIH, CycC-CDK8, and human Taf8-Taf10, complexes were purified following the same protocol as well. Briefly, the cells from 200 ml culture were lysed in 50 ml of lysis buffer: 50 mM Hepes-KOH pH7.6, 400 mM potassium chloride, 10% Glycerol, and 5 mM β-mercaptoethanol, and 0.5 ml of 100 x protease inhibitor mix (6 mM leupeptin, 0.2 mM pepstatin A, 20 mM benzamidine, and 10 mM PMSF). Cell lysate was stirred for 30 min at 4°C followed by centrifugation at 100,000 g for 30 min at 4°C. The supernatant was loaded onto 0.5 ml of Ni-NTA (HIS-Select, Sigma-Aldrich). The resin was washed with 20 ml of the high salt buffer (50 mM Hepes-KOH pH7.6, 1 M potassium chloride, 10% Glycerol, 20 mM Imidazole, and 5 mM β-mercaptoethanol) followed by 5 ml of the low salt buffer (50 mM Hepes-KOH pH7.6, 100 mM potassium acetate, 10% Glycerol, 20 mM Imidazole, and 5 mM βmercaptoethanol). Protein complexes were eluted from Ni-resin with the elution buffer (50 mM Hepes-KOH pH7.6, 100 mM potassium acetate, 10% Glycerol, 300 mM Imidazole, and 5 mM βmercaptoethanol). The recombinant yeast TFIIF complex was purified via an IgG affinity column. Lysate from 200 ml culture prepared as described above, was loaded onto 0.5 ml of IgG resin (GE Healthcare). After wash with the high salt and low salt buffer, the recombinant yeast TFIIF was recovered by TEV protease digestion for 1 hour at room temperature. Concentrations of purified protein complexes were measured by Bradford assay.

### Assay for solubility of the Head module subunits by quantitative western blotting

Cells infected with the various eMOIs were expressed in 50 ml culture of Hi5 cells (1.0 × 10^6^ cells/ml), following the same experimental procedure described above. Cells were harvested in 1.5 ml tubes, cell pellets were frozen in liquid nitrogen, and stored at −80 °C until use. Cell pellets were thawed and were resuspended in 100 µl lysis buffer. Lysates were centrifuged at 15,000 rpm for 10 minutes, at 4°C: supernatant and pellet were separated. Supernatants were mixed with 30 µl of 4x NuPAGE loading buffer (Life Technologies). Pellets were mixed with 120 µl of 1.5 × NuPAGE loading buffer and solubilized by brief sonication. Samples were subjected to 4-12% NuPAGE (Invitrogen), transferred to PROTRAN membranes (Schleicher & Schuell), and probed for the Head module with anti–His tag monoclonal antibody (Thermo Scientific Pierce) for 10xHis-tagged Med17, anti-Med18 (anti-Srb5), and anti-Med11[46]. Detection was carried out using Dylight 680 goat anti–rabbit IgG (Thermo Scientific Pierce) for Med18 and Med11, and Dylight 800 goat anti-mouse IgG (Thermo Scientific Pierce) for 10-His tag on Med17 and scanning with an Odyssey infrared imaging system (LI-COR Biosciences). Bands were quantified using ImageJ software package [47]. 9 pmol (2 µg) of the purified Mediator Head module was used as standard.

## Results and Discussion

### The initial infectivity strongly correlates with overall protein complex production

In our previous work, we used the MultiBac baculovirus expression system as our method of choice for high-yield, recombinant protein production [16]: As illustrated in Figure 1A, the multi-gene construct harboring all 7 genes encoding the Mediator Head subunits was generated, and integrated into baculovirus genome (DH10MultiBac) followed by virus production, expression and purification of the complex [16] (Figure 1A). We often encountered variability in production of the Mediator Head module when switching from one batch of baculovirus to the other, ranging from 3 mg per 1 L culture to 6 mg per 1 L culture (see below). As described in the introduction, given a lack of clarity regarding optimization of protein expression in BEVS, we were prompted to investigate what key factor, if any, is influencing expression of the Mediator Head module. We compared two different batches of baculoviruses harboring the genes for expression of the Mediator Head module: batch MB33 and batch MB88, which exhibited different expression levels: 4.3 mg per 1 L culture from batch MB33, and 6.4 mg per 1 L culture from batch MB88 (Figure 1B). We closely monitored both cell cultures every 24 hours by inspecting cell density and cell morphology. Typically, 24 hours after addition of baculovirus, there was already a marked difference between the two batches: as for batch MB88, the cell density was almost the same as the original cell density of 1.0 x 10^6^ cells/ml, indicating growth arrest (Figure 1C). In addition, the majority of the cells showed rounded cell morphology, slightly increased overall size, as well as expansion of the nucleus, indicative of viral infection. The infectivity of batch MB88 at 24 hours appeared to be 100%. In contrast, after 24 hours the cells infected with batch MB33 virus had a cell density of 1.9 x 10^6^ cells/ml, almost double of the initial cell density. Roughly 50% of cells appeared to be infected judged by their morphologies. The trend continued over the next time points, cells infected with batch MB88 showing 100 % infectivity after 24h, remained unchanged in their cell density until the end of the incubation. (Figure 1C). In contrast, infection with batch MB33 virus was incomplete after 24 hours, and uninfected cells continued to divide until 48 hours (Figure 1C), resulting in a noticeable difference in amount of the cells being harvested (illustrated in Figure 1B below). In short, the discrepancy in the overall expression levels of the Mediator Head module correlates with a difference in initial infectivity of the two batches of virus. Therefore, we hypothesize that the initial infectivity of the baculovirus, which we define here as infectivity 24 hours after virus addition: I_24_, could be an excellent indicator of an overall production of the Mediator Head module.

**Figure 1.**
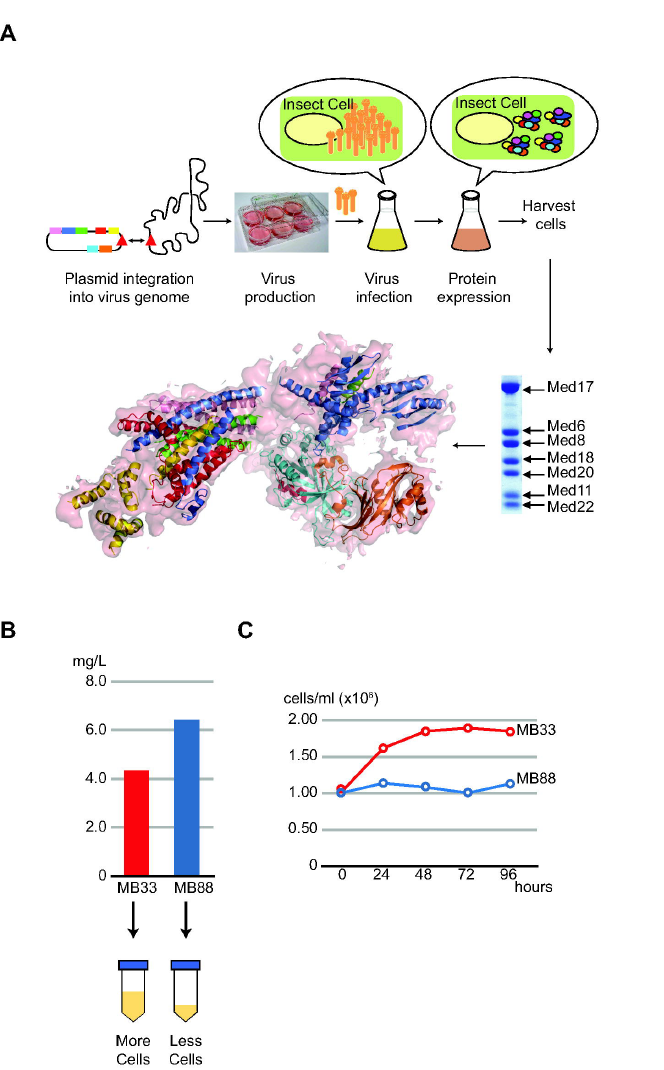
Discrepancy in expression levels of the Mediator Head module was observed when using two different batches of the recombinant baculoviruses, prompting our investigation to seek a key indicator. (A) Schematic diagram of the workflow of the MultiBac baculovirus expression vector system (BEVS) for expression of the Mediator Head module. The single transfer vector harboring genes encoding subunits of the Mediator Head module was integrated into a baculovirus gene followed by virus production, expression and purification of the complex (top). SDS-PAGE of purified Mediator Head module is shown (bottom on right). The structure using the recombinant Head module was determined by X-ray crystallography (PDBID: 3RJ1) (bottom on left). Med17 (blue), Med11 (purple), Med22 (green), Med6 (yellow), Med8 (red), Med18 (cyan), and Med 20 (orange) with electron density map (light pink) (B) Quantity of purified Mediator Head Module varies notably for different batches of the expression baculovirus (top). Batch MB33 in red and MB88 in dark blue (C) Time course for cell density infected by batch MB33 and batch MB88 viruses during expression from time 0 to up to 96 hours.

To test our hypothesis, we pursued to develop a new and simple approach to determine initial infectivitiy (I_24_) in order to examine the correlation between initial infectivities and expression levels of the Mediator Head module in-depth. Technically, flow cytometric assays [35], or the measurement of cell diameter change of post infected cells using Vi-CELL XR Cell Counter (Beckman Coulter) [36], could be used to measure an initial infectivity (I_24_). However, as mentioned above, our goal was to devise an easy and affordable experimental method without the use of specialized equipment and setting so that many end users can utilize it. We reasoned that an initial infectivity (I_24_) could be “estimated” by measuring cell densities: C_0_ (number of cells per ml) being the initial cell density and C_1_ being the cell density at 24 hours after addition of virus, based on the assumption that (i) infected cells lead to immediate growth arrest [48], resulting in no cell division, and that (ii) uninfected cells divide with a growth constant, α in 24 hours: α indicates how much fold change of uninfected cell growth for a 24 hours period, and it is defined as a ratio of cell densities at the time t and at the time, (t+24 hours): α = cell density at (t+24)/cell density at t. For example, α=2 means that cell number doubles in every 24 hours. We like to emphasize at this point, that we were not detecting the number of infected cells directly. Rather we attempted to estimate how many cells are infected based on the consequence of virus infection (infected cells led to cell arrest). Thus, we designate this value as estimate of I_24_: eI_24_. Based on the assumption, the cell density at 24 hours after addition of virus, C_1_ can be described as a sum of (i) the number of infected cells (C_0_ × eI_24_) and (ii) the number of the uninfected cells that were divided [(C_0_ − C_0_ × eI_24_) × α]: C_1_ = C_0_ × eI_24_ + (C_0_ − C_0_ × eI_24_) × α. Therefore, eI_24_ can be described as follows:

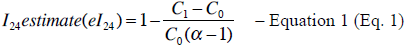

We evaluated if our new method provides estimates of infectivity, that are in line with those measured by commercially available FCA [35]. In this assay, Sf9 cells were infected by serially 10-fold diluted virus stock, and after 24 hours, the number of viral protein gp64 expressing cells were identified by flow cytometry. The initial infectivity can be determined by calculating the ratio between the number of gp64 expressing cells and total number of cells. In parallel, we set up our measurements using Sf9 cells under equivalent conditions in terms of the ratio between virus amount and cell number. The cell densities were measured and the initial infectivity for each virus amount was calculated using our formula (Eq. 1). Clearly, both methods are in good agreement for assessing the initial infectivity at almost all dilution points (Figure 2A), pointing to validity of our newly developed method.

**Figure 2.**
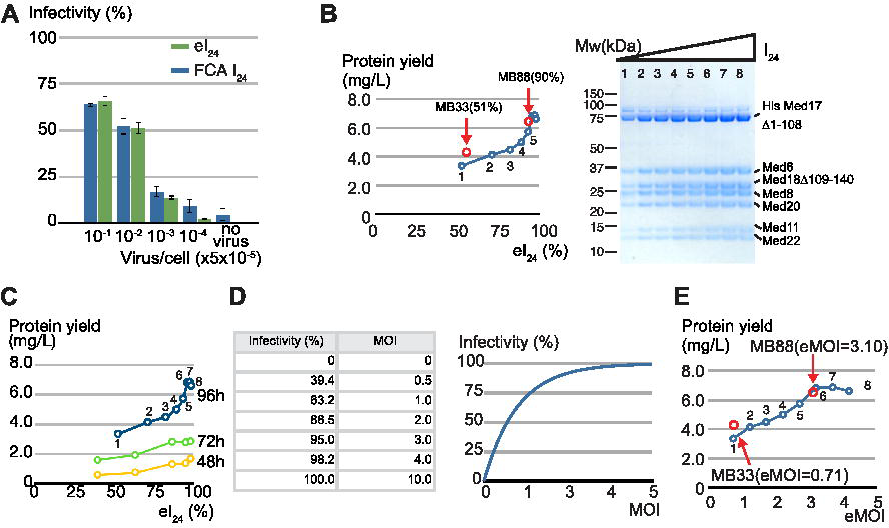
Infectivity of insect cells 24 hours after addition of virus could serve as a good indicator for the expression level of the Mediator Head module. (A) Comparison between the initial infectivity estimates (eI_24_) derived from the cell density measurement based method and I_24_ (FCA) from flow cytometric assay (FCA). Sf9 cells were infected with 10-fold serially diluted recombinant baculovirus expressing the Mediator Head module as indicted in Figure 2A. 24 hours after addition of the virus, infectivity of each dilution point was determined by our cell density measurement based method as well as by FCA (Expression Systems Inc.) in which the percentage of gp64 expressing cells was calculated. X-axis: Virus volume/ total cell number (V_virus_/cell × 5.0 × 10^−5^); Y-axis: initial infectivity (%). eI_24_ are shown in dark blue and FCA I_24_ are shown in green. (B) Initial infectivity (I_24_) vs. yield of the Mediator Head Module. A total of 8 × 200 ml cultures were infected by various amounts of the baculovirus harboring the genes that encode the Mediator Head module. SDS-PAGE of purified Mediator Head module in eight different infectivity conditions is displayed on the right. The protein complex yield from each condition was measured and plotted against eight different infectivities on the left. The estimated infectivity was ranging from 45% to 98.1%. The cell culture conditions of batch MB33 and batch MB88 were superimposed onto Figure 2B indicated by red closed circles and red arrows. (C) Correlation between eI_24_ and the protein complex yield when infected cells were incubated for 48, 72, and 96 hours. A total of 5 × 200 ml cultures were infected by various amounts of the baculovirus harboring the genes that encode the Mediator Head module, and incubated for shorter time periods (48 and 72 hours). The protein complex yield at each infectivity from the culture incubated for 48 or 72 hours was measured, and plotted against five different infectivities. The data from the culture incubated for 96 hours shown in Figure 2B, was superimposed in the same plot. (D) Relationship between infectivity and MOI. Non-linear nature of mathematical relationship between infectivity and multiplicity of infection (MOI) is displayed as table on the left and graph on the right. (E) eMOI vs. yield of the Mediator Head Module. Yield of the Mediator Head Module was plotted against eMOI, which was calculated from eI_24_ values using Eq.4. The cell culture conditions of batch MB33, and batch MB88 were superimposed onto Figure 2E, indicated by red closed circles and red arrows.

With this new simple method in hands, we set up a series of the insect cell cultures with different amounts of the baculovirus expressing the Mediator Head module. Using our formula (Eq. 1), we calculated eI_24_ for each culture and plotted those values against protein yield of the Head module purified from each culture (Figure 2B). In this experiment, infected cells were incubated for 96 hours (our default setting). As shown in Figure 2B, there is a clear correlation between estimated initial infectivity (eI_24_), and protein amount of the Mediator Head module: the Mediator Head module yield increases nearly proportionately to an increase in the value of eI_24_ until it reaches close to 100%. Furthermore, we examined the correlation between eI_24_ and the protein complex yield when infected cells were incubated for a shorter time (48 and 72 hours) (Figure 2C). The length of incubation period influenced total yield of the complex. However, it did not influence the overall pattern of correlation between eI_24_ and yield (Figure 2C). Taken together, these data supports our hypothesis that our estimated initial infectivity, eI_24_, could be a good indicator of the overall Mediator Head module expression in insect cells. The significance of this finding is that despite the complexity of expressing seven proteins simultaneously, the overall protein yield could be influenced by a ‘single’ factor, an initial infectivity, which can be easily estimated by counting cell density using a hemocytometer or cell counter.

### Initial infectivity (I_24_) of a recombinant baculovirus can be mathematically described as a function of multiplicity of infection (MOI), connecting MOI or titer to overall protein complex production

We attempted to mathematically describe initial infectivity, I_24_, as a function of MOI by applying the mathematical formula first presented by Ellis and Delbruck in 1939 [49]: A virus infection is a stochastic event and the probability that number (n) of virus particles infected a cell with a given multiplicity of infection (MOI) can be approximated using Poisson distribution [49], 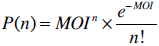, where P (n) is the probability that an insect cell will get infected by number (n) of infectious units 1 baculoviruses. This equation is particularly applicable to our case since an initial infectivity (I_24_) was determined by a primary infection and is not influenced by secondary or tertiary infections. Infectivity as the average percentage of the insect cells that will become infected after addition of baculovirus with a given MOI can be described as: Infectivity = P (n > 0) = 1 − P (0) and P (0) = e^−^MOI.

Therefore, initial infectivity (I_24_) can be described as a function of multiplicity of infection (MOI) as follows:

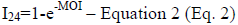

Key findings of this equation are two fold: first, a relationship between I_24_ and MOI is nonlinear (Figure. 2D); and second, more importantly, this equation links MOI directly to overall level of protein expression. Realizing this mathematical relationship between MOI and overall protein complex expression level is key discovery in this work.

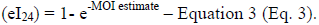

By combining equations 1 and 3, I_24_ estimate = eI_24_, and MOI estimate (eMOI) can be described as follows:

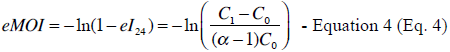

Since MOI = titer/ cell density × cell culture volume (V_c_) × virus volume (V_virus_), then titer estimate, eTiter, can be described as follows:

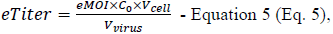
where V_virus_ is a virus volume added to the insect cell culture.

We empirically tested our mathematical formula by utilizing the batch MB33 and batch MB88 viruses, which resulted in different Head module expression levels when infected with an equal volume of virus. The eI_24_ and the corresponding eMOI values of batch MB33 and batch MB88 were determined to be: eI_24_ (batch MB33) = 0.51 (51%), eMOI (batch MB33) = 1.32, and I_24_ (Batch MB88) = 0.90 (90%), eMOI (batch MB88) = 3.1, respectively. Numbers of batch MB33, and batch MB88 and their Head module yields are nicely superimposed on the plot as a function of eI_24_ (Figure. 2B), as well as that of eMOI (Figure. 2E), indicating the validity of our mathematical formula.

Our results so far led us to hypothesize the following as illustrated in Figure 3: first, a single factor, the initial infectivity (I_24_) is a good indicator of overall protein complex expression level as is MOI despite of complexity of multiple proteins being expressed; second, the relationship between I_24_ and MOI is nonlinear (Figure 2D); third, I_24_ could be estimated simply by measuring cell density 24 hours after addition of baculovirus with Eq. 1; fourth, baculovirus “titer” can be estimated using Eq. 5; and finally and most importantly, using eI_24_ or eMOI value of a given virus harboring multiple genes, expression of a protein complex can be quantified and thus, optimized (Figure 3).

**Figure 3.**
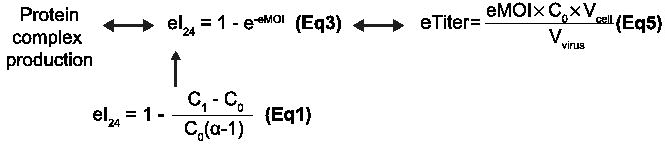
Schematic diagram for our hypothesis illustrating the mathematical relationship among protein production, eI_24_, eMOI, and eTiter. The relationship among protein production, eI_24_, eMOI, and eTiter was displayed as schematic diagram.

### The cell density measurement based method provides estimations of titer values that are in line with those measured by other titer measurement method

Our findings so far were very encouraging in terms of developing a simple and affordable method to estimate viral titer, MOI and infectivity. It will allow us quality control of protein complex production using the BEVS in insect cells at very early stages of the experiment. Moreover, the results from our cell density measurement based method are in good agreement with those from FCA method for assessing the initial infectivity of the baculovirus expressing the Mediator Head module. We further evaluated whether our new method can also provide titer estimates that are in line with those measured by one of the conventional titer measurement assays. To this end, we compared titer values obtained from our formula (Eq. 5) (Figure 3) to those from commercially available flow cytometric assay based titer measurements, which provides titer values, that are in good correlation with those obtained by plaque assay [35]. Titers of a total of twelve baculoviruses were measured by both methods and compared (Figure 4A and 4B). Six of these virus do not contain any particular gene and consist of the control virus with known titer [1.0 × 10^9^ Infectious Unit (IU) /ml] purchased from Expression Systems, Inc (Davis, CA), baculovirusgenerated from bacmid alone (DH10MultiBac) (Supplemental Figure S1A) [23], eYFP containing bacmid [50] (Supplemental Figure S1B), bacmids fused with several empty MultiBac transfer vectors, one vector (pFL) labeled as Vec1, (Supplemental Figure S1C), two vectors ( pFL+pUCDM) labeled as Vec2, (Supplemental Figure S1D), or three vectors (pFL+pUCDM+pSPL) labeled as Vec3, (Supplemental Figure S1E) (Figure 4A). The other six baculoviruses contain genes for the protein complexes, which include the Mediator Head module, yeast TFIIF, human Taf8-Taf10 heterodimer, yeast CycC-CDK8 complex, yeast core TFIIH, and the Mediator Middle module, respectively (Figure 4B). As described in the supplemental protocol, we derived eTiter values from four different data points of different virus volumes using linear regression: Each measurement was repeated three times and averaged (Supplemental Figures S2-3). Comparing both methods for the viruses expressing protein complexes, the obtained titer values are in good agreement for five out of six viruses (Figure 4B). The other viruses, which do not contain genes for recombinant protein expression, and the one encoding the yeast core TFIIH genes titer values measured using our formula (Eq. 5) are on average 2.1 times higher than those determined by the FCA method. In this regard, there are several other studies published comparing different titer measurement methods. For example, a 2-3 fold difference between the end-point dilution assay, and qPCR based assay was observed [51]; the analysis of end point dilution using GFP and anti-DBP (DNA Binding Protein) based assay showed as much as a 15-fold difference between the titer values [52]. Compared to these previous results, clearly, our measurement results are in a reasonable and acceptable range. Therefore, we conclude that our formula provides good titer estimations and thus, can be extended in its application to aid in the robust expression of many proteins.

**Figure 4.**
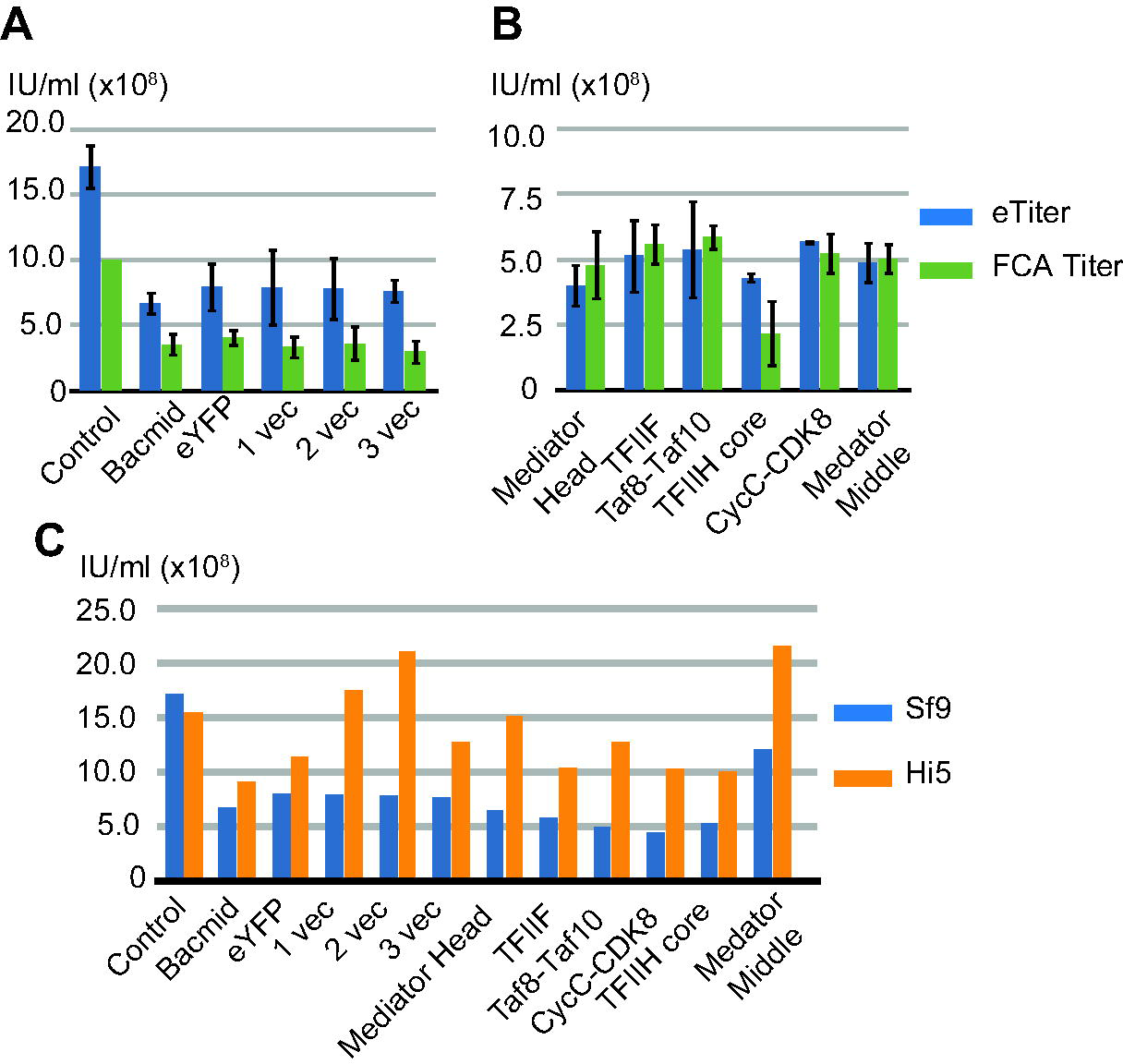
Comparison of titer values by the cell density measurement based method vs. commercially available FCA based titer measurement. (A) Titer values were obtained from the cell density measurement based method and commercially available flow cytometric assay (FCA) based titer measurement for a total of twelve baculoviruses, and these values were compared. All measurements were performed in Sf9 cells. Titers of the control virus with known titer (1.0 × 10^9^ IU/ml) purchased from Expression Systems, baculoviruses generated from bacmid alone [23], eYFP containing bacmid [50], bacmids fused with several empty Multibac transfer vectors, respectively. Each measurement was repeated three times and averaged except for the control virus. eTiters are shown in dark blue and FCA titers are shown in green. (B) Comparison between eTiter and FCA titer. Titers of recombinant baculoviruses for the yeast Mediator Head module, yeast TFIIF, human Taf8-Taf10 heterodimer, yeast CycC-CDK8 complex, yeast core TFIIH, and the yeast Mediator Middle module were measured by two different methods and compared. Each measurement was repeated three times and averaged. eTiters are shown in dark blue and FCA titers are shown in green. (C) Different insect cell lines yield different eTiter values. The eTiters for a total of 12 viruses were measured in both Sf9 and Hi5 cells and compared. The eTiters in Sf9 are colored in dark blue, and those in Hi5 are colored in orange.

As suggested, titer values can be dependent upon medium and cell types [35]. To address this, we tested if there is a difference in eTiter values between Sf9 cells and Hi5 cells. Since the FCA titer measurement service is available in only Sf9 cells, but not in Hi5 cells, we used our method to measure eTiter values of all twelve viruses described above using Hi5 cells and compared them to Sf9 cells (Figure 4C, Supplemental Figures S4). Although eTiter values of the control virus are similar in both cell lines, eTiter values of the other eleven MultiBac based viruses measured in Hi5 cells are approximately twice as high than those measured in Sf9 cells.

### The condition at which eI_24_ ≈100% (eMOI=3 or greater), provides the optimal or near optimal expression levels for a series of protein complexes involved in eukaryotic transcription by RNA polymerase II

We applied our newly developed method to several other recombinant protein complexes involved in eukaryotic RNA polymerase II (Pol II) mediated transcription to see if their expression levels could be quantified and optimized in a similar fashion. Eukaryotic transcriptio by Pol II is driven by a series of multi-protein complexes including the TATA-box protein associated factors (TAFs), general transcription factors, TFIIH, and TFIIF, Mediator, and CDK8 module [53–55]. Generation of recombinant forms of these complexes and their sub-complexes will be essential for structural and functional studies. A set of expression baculoviruses for human Taf8-Taf10 complex (2 subunits)[43], yeast core TFIIH (6 subunits), yeast TFIIF (3 subunits), yeast CDK8-CycC (2 subunits), yeast Mediator Middle module (8 subunits), and the yeast Mediator Head module (7 subunits) were generated. Following our protocol (Supplemental Protocol), eI_24_ value of each virus was determined followed by calculating eMOI, and eTiter (Figure 5). These complexes were expressed in 8-9 different infection conditions ranging from 18% infectivity (eMOI=0.2) to ~100% (eMOI=3-5) (Figure 5). Each complex at each infection point was expressed in a 200 ml culture of Sf9 or Hi5 cells followed by affinity purification and quantification of the purified complex. The yield of each protein complex was plotted against eI_24_. We also used the corresponding eMOI to display because eMOI is intuitively easy to understand: for instance, eMOI=2 indicates adding twice as much volume of virus as eMOI=1. There is a good correlation between the eI_24_ of the expression baculovirus and the overall protein complex production (Figure 5). Protein complex yield peaked at or near infection saturation point of eMOI being 3 or greater (Figure 5) except for the case of the Mediator Middle module: its yield peaked at an earlier infectivity point (39.3%; eMOI=0.5) (Figure 5A). However, the protein complex yield difference between eMOI=0.5 and eMOI=3-5 is relatively small with 10 mg/L and 9 mg/L, respectively, accounting for only a 10% difference. In other words, the eMOI=3-5 is nearly optimal. Key conclusion is that the condition at which eI_24_ =100% or eMOI≥3 provides the optimal, or near optimal expression levels for all cases tested. We call this the “eMOI=3 or greater” rule. It is not entirely surprising to see that optimal protein expression is observed when cells are fully infected within the first 24 hours, considering a delay in infection due to lower virus load consequently postpones reaching the level of maximum amount of recombinant protein producing cells and ultimately yield in the given time-frame. Our “eMOI=3 or greater” captures such condition for an optimal protein production.

**Figure 5.**
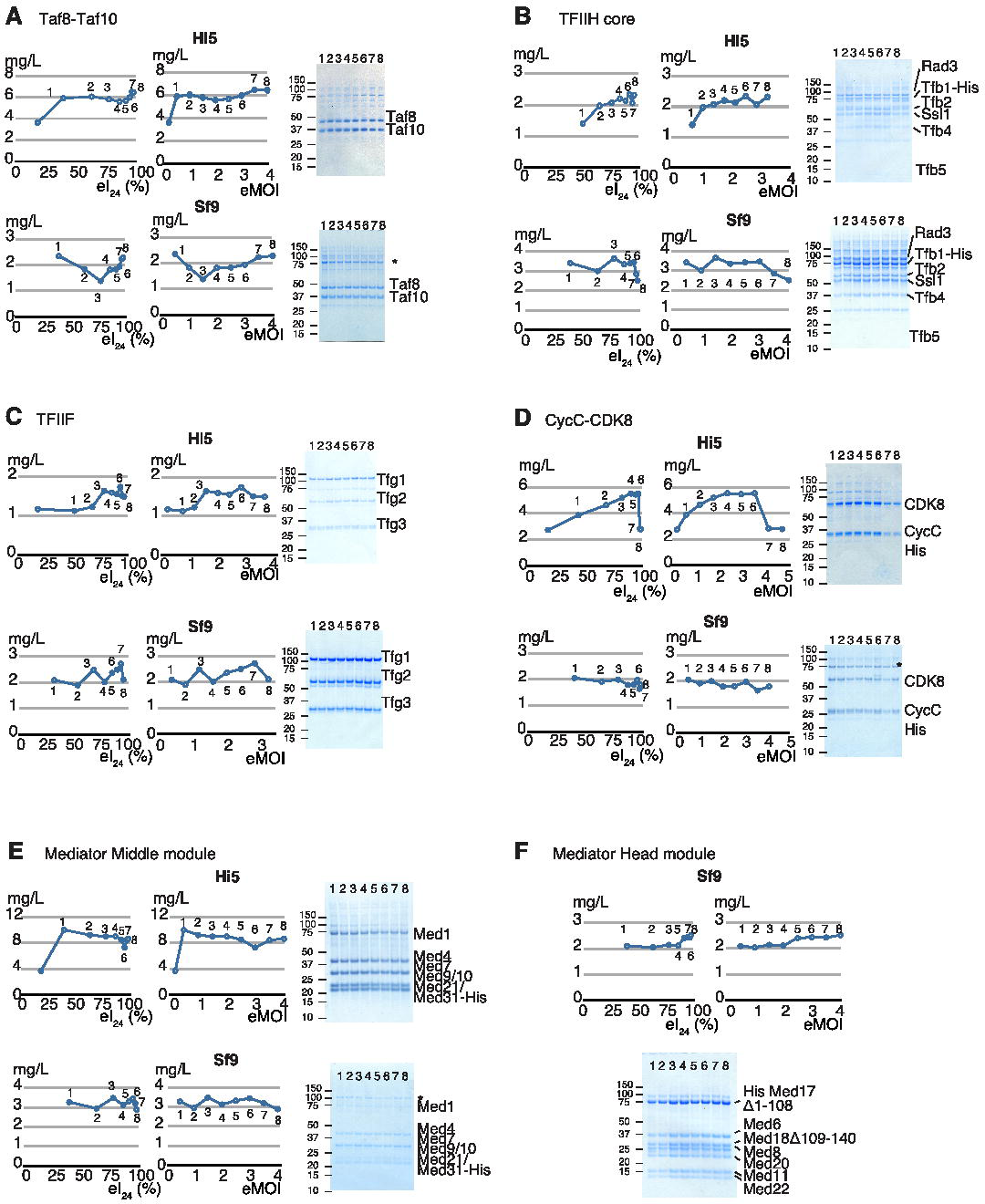
Quantification of expression of the multi-protein complexes involved in eukaryotic RNA polymerase II transcription by varying eI_24_ or eMOI values in the Hi5 and Sf9 cells. A total of six different multi-protein complexes were expressed in Hi5 cells and Sf9 cells under different conditions in terms of initial infectivity (eI_24_) or eMOI ranging from eI_24_ = 18 % (eMOI=0.2) to eI_24_=100% (eMOI=4-5) in a 200 ml culture scale. SDS-PAGE of each purified complex with different eI_24_ or eMOI values is shown in right. Yield of complex at each point was measured and plotted against I_24_ (left) or eMOI (right) panel. The results from expression using Hi5 cells were shown on top and those from Sf9 cells were shown in the bottom. (A) Human Taf8-Taf10 complex, (B) yeast core TFIIH complex, (C) yeast TFIIF, (D) yeast CycC-CDK8, (E) the Mediator middle module, and (F) the Mediator Head module. Asterisk (*): contaminant

We further tested the idea that once the cells are more than 95 % infected (eMOI ≥ 3), any additional excess virus would not necessarily increase overall expression level of protein complexes. To test this idea, the Mediator Head module, hTaf8-Taf10 and yeast core TFIIH were chosen because the Head is our model complex and other two are on different ends of the spectrum in size and expression levels: hTaf8-Taf10 is considered a relatively small complex (81 kDa), and was highly expressed (Figure 5A) while yeast core TFIIH is a relatively large complex (6 subunits, 320 kDa) and not expressed well (Figure 5B). For all three complexes (Supplemental Figure S5), additional excess virus did not improve overall yield of the protein complexes, suggesting that once the cells are fully infected, further addition of expression virus may not improve overall expression of protein complexes. During our study, we noticed a marked difference in expression levels of protein complexes between Hi5 and Sf9 cells. Performance of Hi5 cells exceeded Sf9 cells in 4 out of 6 complexes (Figure 5A, 5D-F). For expression of yeast core TFIIH (Figure 5B) and TFIIF (Figure 5C), Sf9 cells performed better than Hi5 cells, suggesting that both cell lines should be tested to obtain maximum yield of protein complex of interest.

Finally, we tested if an increase in yield of protein complex could be attributed to an increase in overall expression of the complex or an increase in solubility of the subunits. We used the Mediator Head module as a test case since it showed most change in its yield between low and high eMOI (Figure 2D). We compared soluble and insoluble fractions of the Head module with different eMOI ranging from eMOI=0.1 (10% infected initially) to eMOI=4 (~100% infected) by immunoblotting using antibodies against Med17, Med18 and Med11 subunits (Supplemental Figure S6A-C). The ratio between soluble and insoluble fractions of Med17, Med18 and Med11 appears to stay constant, suggesting that an increase of I_24_ or eMOI leads to an increase in overall production of the Mediator Head subunits.

The results of a total of six multi-protein complexes are consistent with our hypothesis that eI_24_(or eMOI) can be an excellent indicator of expression levels of multi-protein complexes. By varying eI_24_ or eMOI value, the optimum expression points of all complexes can be identified. But in the end, the condition at which eI_24_ =100% or eMOI ≥3 provides the optimal, or near optimal expression levels (“eMOI=3 or greater” rule). In essence, our simple and affordable cell density measurement based method provides eI_24_ or eMOI value of a recombinant baculovirus. Researchers (end users) can use these values to quantitatively optimize expression levels of proteins or protein complexes of their interest. We named our newly developed quantitative method “TEQC” method: “Titer Estimation for Quality Control” (TEQC) of protein or protein complex production.

### TEQC method can be utilized to achieve reproducible expression of protein complexes to ensure quality control

We tested whether we could use the eMOI value to achieve reproducible expression of protein complexes, resulting in a production of the recombinant protein complexes with maximum yield consistently. The conditions for maximum expression levels of 6 protein complexes are as follows: in the Hi5 cells, the Mediator Head module peaked at eMOI of 3.7; hTaf8-Taf10 at eMOI of 3.5, yeast CDK8-CycC at eMOI = 3.5, and Mediator middle module at eMOI= 0.5; in SF9 cells, yeast core TFIIH at eMOI of 3.0, and yeast TFIIF at eMOI of 3.0 (Figure 5), respectively. For simplicity, we are referring to eMOI only. All these six protein complexes were expressed using the same optimum eMOI and cell line but were expressed on different days. These independent expressions were repeated three times for all complexes. The yield of each purified protein complex was measured, the numbers were averaged, and compared to the original expression level that provided the optimum eMOI value in that cell line. For all tested complexes, the yield of purified protein was highly reproducible (Figure 6A).

**Figure 6.**
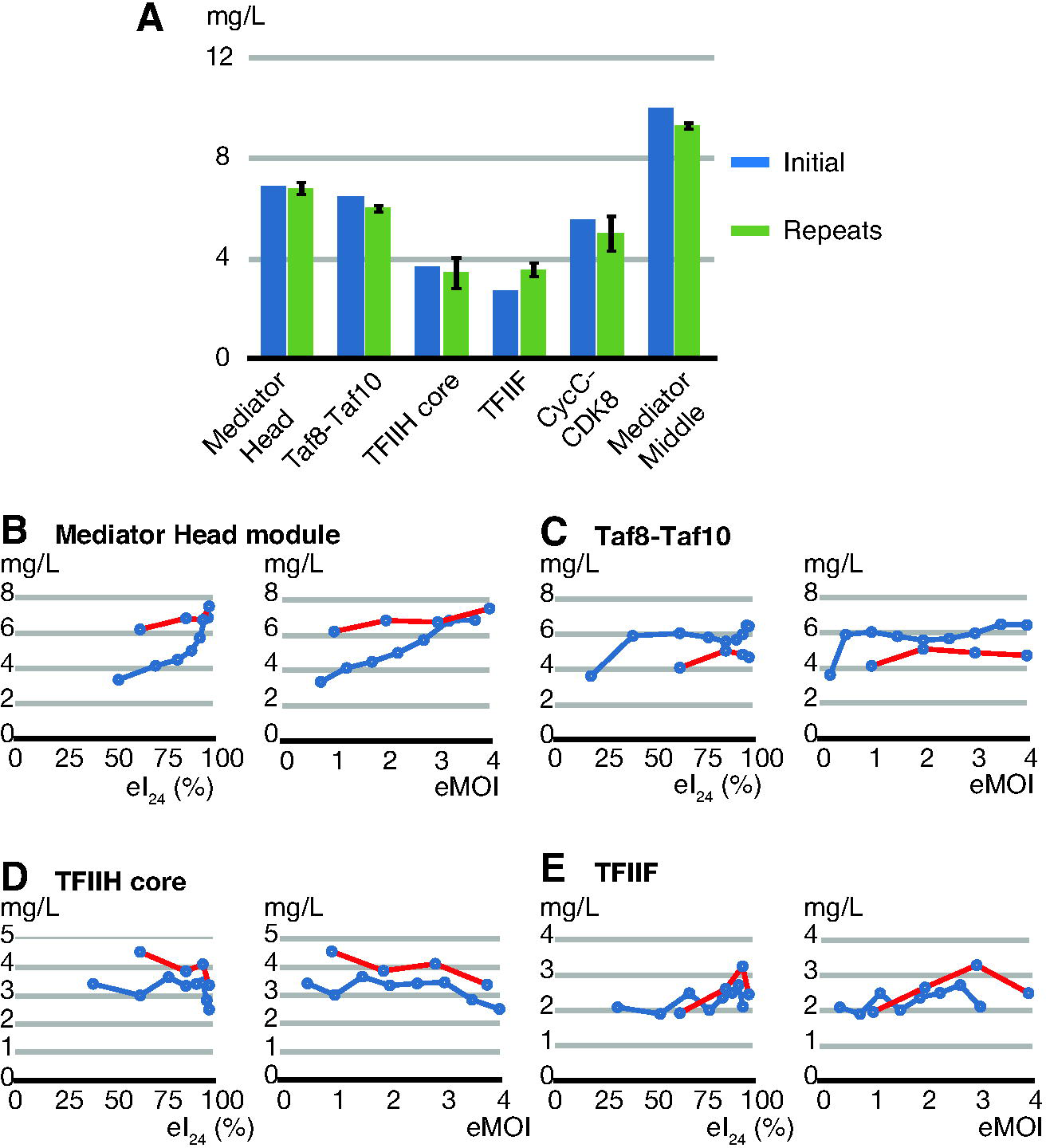
TEQC method ensures reproducibility of production of the multi-protein complexes. (A) Mediator Head module, Taf8-Taf10, TFIIH core, TFIIF, CycC-CDK8, and Middle were expressed at their optimum infectivity or eMOI, and each protein complex yield was measured; labeled as “initial” (dark blue). Expression was repeated three times using the same eMOI or infectivity value for each complex independently, purified protein complex yields were measured and the numbers were averaged; labeled as repeats. The repeats (green) were compared with the initial protein expression optimization results (dark blue). (B-E) Mediator Head, Taf8-Taf10, core TFIIH, and TFIIF complexes were expressed using freshly made new high titer viruses as well as the old viruses with declined titers. The eTiter values of both new and old viruses were determined and volume of each old or new virus was adjusted such that the expression of each complex was carried out at the same eI_24_ or eMOI. After expression and purification, protein complex yield was measured, and the data were plotted against either eI_24_ or eMOI. (B) Mediator Head module was expressed in Hi5 cells (red) (C) Taf8-Taf10 was expressed in Hi5 cells, (D) TFIIH core was expressed in Sf9 cells, (E) TFIIF was expressed in Sf9 cells, and compared with the initial expression results (blue) and the results from old viruses (red).

One of the BEVS benefits is that once a virus is made, it is stable for a long period of time, ranging from up to couple of years if stored at 4 °C to much longer time spans for frozen stocks, allowing it be used for multiple rounds of infections and experiments. Over time though, the infectivity of the virus decreases. To address the situation in which one would like to use expression baculovirus that has declined titer due to prolonged storage at 4 °C we hypothesize as follows: as long as the eI_24_ or eMOI value is the same, an old virus with declined titer could maintain the same level of expression of protein complex as when it was newly made. To test this, we set up expressions of four protein complexes described above with eMOI from 1.0 to 4.0, using the old viruses that had been kept at 4 °C for an extended period of time (8 to 10 months). Their eTiters had been declined as follows: Mediator Head from 7.3 × 10^8^ to 4.0 × 10^8^ (IU/ml), hTaf8-Taf10 1.2 × 10^9^ (IU/ml) from to 6.7 × 10^8^ (IU/ml), core TFIIH from 4.5 × 10^8^ to 3.3 × 10^8^ (IU/ml), and TFIIF from 5.1 × 10^8^ to 3.8 × 10^8^ (IU/ml), respectively. eMOI values ranging from 1 to 4 were achieved by increasing the volume of each virus to compensate for the loss of virus titer. The yield of each purified protein complex was measured and compared to the original expression level (Figure 6B-E, blue lines) by superimposing the data onto the plots of the original expression data indicated by red lines (Figure 6B-E). Overall, when the eI_24_ or eMOIs were adjusted, the expression level profiles of the protein complexes generated by the old viruses were similar to those generated by their original viruses (Figure 6B-E), strongly suggesting that eI_24_ or eMOI is the key indicator of expression level of protein complexes.

## Conclusion

This work has revealed the connection among initial infectivity, MOI, and an overall recombinant protein production. We developed a simple and affordable cell density measurement based method to estimate initial infectivity, MOI, and titer, leading toward the development of the TEQC method for optimization of a production of protein complexes using the BEVS. Key finding is that optimal or near optimal expression levels are reached when eMOI ≥ 3, which our “eMOI=3 or greater” rule. Since expression of single proteins could be viewed as “one” subunit multi-protein complex, the TEQC method should be applicable to expression of single proteins as well. Additionally, expression of a protein complex in insect cells was carried out in this work by infecting a single virus harboring multiple genes encoding the subunits of the complex, and not by co-infecting multiple viruses. The strategy for optimization of protein complex expression using the co-infection method will be a future subject of investigation.

## Author contributions

T. I. and Y. T. conceived the idea and designed experiments; T. I., M.B., K.Y., S. W., performed the experiments. S.W. T. I. and Y. T. analyzed the data; Y. T. supervised the study; and T. I., S.W., and Y.T. prepared figures and wrote the paper.

## Acknowledgements

We thank Dr. B. Hayes for providing the raw FCA data and technical help. We thank Drs. H. Hundley, I. Berger and S. Trowitzsch for critical reading of the manuscript. We thank Drs. I. Berger and S. Trowitzsch for providing human Taf8-Taf10 construct and Dr. F. Ponticelli for providing pDt/g1g2 vector. We thank Dr. H. Noguchi for his help generating a part of the data. This research was supported by US National Science Foundation grant MCB-1157688, the National Institutes of Health (R01 GM111695), and Showalter Trust Fund to (Y.T.).

